# Evidence for a Transient State of Auditory Hypersensitivity During Initial Onset of Tinnitus, Evidenced by Intensity Dependence of the Auditory Evoked Potential (IDAEP)

**DOI:** 10.1101/2025.08.02.668306

**Authors:** Abishek Umashankar, William Sedley, Kai Alter, Phillip E. Gander

## Abstract

Our understanding of tinnitus pathophysiology may be greatly advanced by understanding how the condition evolves from its initial onset or acute stage to its chronic manifestation. Such a transition likely reflects dynamic neurophysiological changes within central auditory and non-auditory networks. Our integrated model of tinnitus posits that sensory precision (sensory weighting) may be heightened during the acute stages of tinnitus to resolve degraded auditory input, but in chronic tinnitus, its role may diminish as plastic processes take over the percept’s maintenance. Consequently, we hypothesize that bottom-up neural mechanisms linked to initiation of tinnitus, such as central gain and neural synchrony, are maximal around the time of tinnitus onset, but later subside by way of regression to the mean. We evaluated this hypothesis by measuring central auditory reactivity through the Intensity Dependence of Auditory Evoked Potential (IDAEP), a non-invasive index of higher-order inhibitory processing within the auditory system. A steeper IDAEP slope is associated with heightened sensory reactivity (higher sensitivity to changes in auditory stimuli), indicative of reduced central inhibition. Conversely, a shallower slope reflects greater inhibitory control. Studying a group with acute tinnitus (onset within six weeks), with a repeated assessment after six months from onset, we found an initially increased IDAEP slope in the acute stage, which had significantly reduced at follow-up, supporting our hypothesis that there is increased sensory reactivity during tinnitus onset, which need not persist in order for tinnitus to become chronic.

## Introduction

Studying acute tinnitus is essential for identifying early biomarkers and mechanisms that differentiate transient from persistent forms or stages. The exact temporal demarcation from acute to chronic tinnitus is not standardised, with arbitrary cut-offs varying between 3 and 12 months (Haider et al., 2018). However it is defined, the acute window might help us understand the neurophysiological and neurochemical processes that first cause tinnitus, hopefully leading to advances that inform the development of targeted therapies and improve prognosis.

Tinnitus is often understood as a disorder of altered central gain (a multiplicative increase in the input-output function), with central auditory pathways exhibiting maladaptive changes in excitability and response dynamics (Auerbach et al., 2014; Zeng, 2013). Our working model of tinnitus based on sensory precision offers a framework for conceptualising the transition from acute to chronic tinnitus. In the acute phase, the brain responds to degraded auditory input such as from hearing loss by upregulating sensory precision and/or gain, effectively increasing the internal weighting of ambiguous auditory signals to resolve uncertainty. This heightened precision can inadvertently amplify internal noise, leading to the false perception of sound. Over time, even if sensory precision returns to baseline, chronic tinnitus may persist due to neuroplastic changes, such as altered predictive coding, memory consolidation, and perceptual learning, which continue to sustain the tinnitus percept in the absence of ongoing peripheral input abnormalities (Sedley et al., 2016).

One type of measurement tapping into tinnitus-related neural reactivity (especially into mechanisms such as central gain) that would help us measure the changes between acute and chronic tinnitus could be the usage of evoked potential measures, as these capture stimulus-driven neural activity. In particular, the N1-P2 complex of the auditory evoked potential has been widely employed in both tinnitus and basic auditory research as an objective index of cortical sound processing. This complex is sensitive to auditory change detection, sensory gating, attentional allocation, and hypervigilance (Mohebbi et al., 2019; Spielmann Moura et al., 2010) with N1 being localized to the superior temporal plane, and P2 having multiple generators in the primary and secondary auditory cortices of both hemispheres (Lightfoot, 2016). However, N1-P2 findings relating to tinnitus have been inconsistent, yielding varied results regarding their magnitude and latency (Azevedo et al., 2020), and it is hard to draw any clear conclusions.

A particularly informative extension of the N1-P2 complex is the Intensity Dependence of Auditory Evoked Potentials (IDAEP), which utilizes the slope of the N1-P2 that quantifies the change in response amplitude per unit increase in stimulus intensity. IDAEP is typically measured at low stimulus frequencies, up to 1 kHz. A steeper IDAEP slope is associated with increased reactivity to sensory input, while a shallower slope reflects stronger inhibitory modulation (Fitzgerald et al., 2009; Zhang et al., 2017). The slope is a key indicator of certain aspects of central auditory gain, as it captures how neural response amplitude changes with increasing stimulus intensity.

In this study, we hypothesised that the acute stage of tinnitus would be characterised by elevated IDAEP slope specifically, and this would reduce as the chronic stage was reached; this was partly motivated by our general hypothesis about transiently increased gain being implicated in initial tinnitus causation (Sedley et al., 2016), and partly from the results of a pilot study in our lab (unpublished data from Sedley et al. (2019)), which did not use an IDAEP paradigm, but nonetheless found greater N1 amplitude differences, between stimuli with a 6 dB intensity difference, in the acute stages of tinnitus compared to after 6 months in the same individuals. As such, we specifically predicted that IDAEP would be increased around the time of tinnitus onset, and then reduce again when entering the chronic stages.

## Methods

### Participants

We studied 29 participants with Acute Tinnitus which we defined as a duration of less than 6 weeks, with mean tinnitus duration 3.93 weeks (SD 2 weeks) and a median duration of 4 weeks with one outlier with a duration of 12 weeks. The outlier subject was included in this study as she fell within the time limit identified by previous authors as acute, specifically a length of less than 3 months (Vielsmeier et al., 2020; Shim et al., 2011), and also participated in the follow-up study.

We also recruited 25 participants with Chronic Tinnitus, defined as a duration more than 6 months, with a mean duration of 8.82 years (SD 8.38 years) and a median duration of 5 years, and 18 non-tinnitus Controls. Participants were recruited through community advertising on Google Ads for Acute and Chronic tinnitus, and internally within Newcastle University’s research volunteer pool for Chronic tinnitus and Control groups. The 29 Acute Tinnitus participants were invited for reassessment after a minimum of 6 months from tinnitus onset which we took to indicate their chronic stage, and 15 of them volunteered for and completed this further testing. We refer to this as the ‘Post-acute’ group, to distinguish it from the group recruited during the chronic stage of tinnitus. Individuals aged 18 and older capable of providing informed consent were included, but those with Meniere’s disease, epilepsy, or middle ear pain or infections were excluded from the study.

Control participants were matched individually to those in the Acute Tinnitus group for age, sex, and hearing. Control participants were presented with stimuli that matched the frequency and presentation ear(s) of their matched tinnitus participants. An attempt to match the Chronic Tinnitus group to the other groups in these regards was only partially successful. Details are displayed in Table 1.

**Table 1:** The participants demographics for each group are shown in. **Table 1**. N denotes number of samples, M denotes mean, SD denotes Standard Deviation, MD denotes Median, F statistic indicates a One Way ANOVA has been carried out, H statistic with X^2^ indicates a non-parametric Kruskal Wallis test, X^2^ indicates a chi square goodness of fit test, NS denotes No Significance. * indicates presence of statistical significance at 95% confidence interval. Hearing thresholds were calculated by averaging the left and right ear values in cases of bilateral tinnitus, whereas for unilateral tinnitus, the threshold was based on the ear with tinnitus.

| **Subjects included in 1 kHz analysis** |  |  |  |  |  |
| --- | --- | --- | --- | --- | --- |
| **Parameters** | **Acute**<br>**Tinnitus** | **Chronic**<br>**Tinnitus** | **Controls** | **Significance**<br>**Overall** | **Post hoc**<br>**Significance** |
| **Number of**<br>**Subjects** | 27 | 20 | 17 |  |  |
| **Age (Years)** | M = 52.89<br>SD = 13.56 | M = 54<br>SD = 9.5 | M = 53.53<br>SD = 15.54 | F = 0.43<br>p = 0.958 | NS |
| **Gender**<br>**(M/F)** | 5/22 | 9/11 | 3/14 | $X^2 = 5.737$<br>p = 0.079 | NS |
| **Ear**<br>**presentation**<br>**(U/L – B/L)** | 12/15 | 7/13 | 6/11 | $X^2 = 0.569$<br>p = 0.752 | NS |
| **Hearing (1**<br>**kHz) dBHL** | M = 12.31<br>SD = 8.20 | M = 15.5<br>SD = 15.87 | M = 10.75<br>SD = 11.48 | F = 0.791<br>p = 0.458 | NS |
| **Subjects included in Tinnitus Frequency analysis** |  |  |  |  |  |
| **Number of**<br>**Subjects** | 25 | 13 | 15 |  |  |
| **Age (Years)** | M = 51.8<br>SD = 13.59 | M = 54.38<br>SD = 9.94 | M = 51.87<br>SD = 15.17 | F = 0.184<br>p = 0.833 | NS |
| **Gender**<br>**(M/F)** | 5/20 | 4/9 | 2/13 | $X^2 = 1.304$<br>p = 0.521 | NS |
| **Ear**<br>**presentation**<br>**(U/L – B/L)** | 10/15 | 5/8 | 5/10 | $X^2 = 0.181$<br>p = 0.913 | NS |
| **Hearing (4**<br>**kHz) dBHL** | M = 20.6<br>SD = 14.83 | M = 25<br>SD = 13.42 | M = 16.33<br>SD = 13.69 | F = 1.301<br>p = 0.281 | NS |
| **Hearing (8**<br>**kHz) dBHL** | M = 25.2<br>SD = 17.73 | M = 461.15<br>SD = 16.15 | M = 28.17<br>SD = 23.87 | F = 3.005<br>p = 0.059 | Acute<br>Tinnitus –<br>Chronic<br>Tinnitus:<br>p = 0.05* |

### Audiological Assessment

Prior to the experiment, participants completed a pre-screening form containing questions designed to confirm eligibility based on the inclusion and exclusion criteria. The experimenter, a qualified clinical audiologist with experience in tinnitus, conducted interviews with the participants prior to the experiment, confirming a medical history consistent with subjective tinnitus and the lack of atypical symptoms or concerns about an unidentified underlying cause. Participants provided demographic information, including age and sex, as well as information about their tinnitus, such as type (tonal, noise-like, or other), duration, which ear(s), and a past physical and mental health history, including any otological disorders. Four common, validated questionnaires—the Tinnitus Handicap Inventory (THI) (Newman et al., 1996), the Tinnitus Functional Index (TFI) (Meikle et al., 2012), the Hyperacusis Questionnaire (HQ) (Khalfa et al., 2002), and the Inventory of Hyperacusis Symptoms (IHS) —were used to evaluate the impact and distress related to tinnitus and any co-existing hyperacusis symptoms (Greenberg & Carlos, 2018).

Participants underwent a Pure Tone Audiometry (PTA) test to establish hearing thresholds at octave frequencies ranging between 250 Hz up to 8 kHz. Thresholds were estimated based on the initial presented level at either 40 dB HL or above depending on the participant’s residual hearing. If perceived audible, the stimulus was continuously reduced by 15 dB HL until the participant did not perceive it as audible. From the first reversal, a 5 dB up, 10 dB down staircase procedure was then used until the final threshold was established by two positive responses out of three trials (Carhart & Jerger, 1959).

### Tinnitometry

Tinnitus pitch and loudness were determined via a *tinnitometry* procedure, for participants with either acute or chronic tinnitus. In cases of unilateral tinnitus, matching stimuli were presented contralaterally, and in bilateral tinnitus they were presented bilaterally. Participants first matched their tinnitus for loudness and then for frequency from a starting reference stimulus of 6 kHz. In bilateral cases where the tinnitus was asymmetric, additional adjustments were made to balance the two ears. Both loudness, then frequency, were iteratively adjusted until the sound closely matched the participant’s tinnitus, and no further changes were judged necessary by the participant. This procedure was repeated across three trials and a mean of the trials was taken and considered as the final loudness and pitch of the tinnitus. The stimulus used for this procedure was either a pure tone or narrow band noise (1/3 octave with Hanning spectrum) depending on the subjective similar bandwidth of the tinnitus.

### IDAEP Stimuli

With a total of 200 stimuli for each of 3 intensities and 2 frequencies, and a 25-minute total experiment length, the IDAEP stimuli consisted of pure tones at 1 kHz and the individual’s matched tinnitus frequency (or, for controls, the matched tinnitus participant’s tinnitus frequency), with a duration of 100 ms, onset/offset ramp of 10 msec, and an interstimulus interval uniformly randomized between 1 sec and 1.5 sec. For each frequency (1 kHz and tinnitus-matched), three different intensities were presented (details of the intensities are explained below). Stimuli were presented in blocks, with each block containing stimuli of a fixed frequency and random intensity, with a total of 60 stimuli per block (i.e. 20 per intensity), and a total of 10 blocks per frequency. To compensate for hearing loss, loudness recruitment, and hyperacusis, stimulus intensities were individually adjusted based on each participant’s dynamic range—the span between hearing threshold and uncomfortable loudness level—for both 1 kHz and the tinnitus frequency. Hearing thresholds were established for the experimental stimuli using an ascending-descending run of 3 dB steps until at least a 50% positive response was obtained (i.e., two correct responses out of four trials) (Carhart & Jerger, 1959). The uncomfortable loudness level was established by presenting an ascending run of 3 dB step size from the threshold until the participant just perceived the tone to be uncomfortably loud or showed signs of discomfort (Carhart & Jerger, 1959). Dynamic range was defined as uncomfortable loudness level (ULL) minus hearing threshold. Stimulus intensities were determined as follows:

a. Low Intensity: hearing threshold plus 60% of Dynamic Range
b. High Intensity: ULL – 10 dB
c. Medium Intensity: Mean of Low and High Intensity

### ULL: Uncomfortable Loudness level, DR: Dynamic Range

For participants with (or controls matched to subjects with) unilateral tinnitus, stimuli were only presented in the tinnitus ear; for those with bilateral tinnitus, they were presented in both ears. The controls also were individualised in intensity based on the same dynamic range procedure. Table 1 shows the demographic details and distribution of presentation ears for each group.

### IDAEP EEG Recording

EEG was recorded using a 64-channel Active Two system (Biosemi) in a soundproof room. Electrode offset was kept at the manufacturer’s recommended limits of ±10 mV with a sampling rate of 256Hz.

The experiment was a passive task, during which the participants watched a silent subtitled movie of their choosing. All stimuli were generated and presented using Matrix Laboratory (Matlab) version R2019a and Psychtoolbox version 3 (Pelli, 1997).

### EEG Preprocessing

Data were processed in Matlab, version R2019a, using the EEGlab toolbox (Delorme & Makeig, 2004) and customised code. Data were re-referenced to the P9/P10 (approximating a linked mastoids montage). Bandpass (non-phase-distorting) filtering was performed between 1 Hz and 30 Hz. Channels judged to contain excessive noise were removed and reconstructed by interpolation from neighbouring channels. Channel rejection was based on visual inspection and a correlation higher than 0.8 to other channels were also rejected to improve signal quality. This was followed by epoching between - 0.1 and 0.5 s from stimulus onset. Independent Component Analysis (ICA) was performed, and components capturing predominantly eye blinks or eye movements were removed from the data. Subsequent to ICA, artifact rejection was also carried out using individual component rejection where components were auto rejected using a probability of 5 and Kurtosis of 8 as the threshold. Baseline correction was performed based on the period of −100 to 0 msec. All trials for each stimulus frequency and intensity were then averaged to yield the event-related potential.

### Data Analysis

After preprocessing, we used customised MATLAB code to perform automatic peak detection to determine the N1 and P2 for each subject. At electrode FCz, we measured the most negative peak amplitude for N1 between post-stimulus latency 90 and 200 msec and the most positive peak amplitude for P2 between post-stimulus latency 100 and 250 msec. The peak-to-peak amplitude between N1 and P2 in μV was taken as the variable of interest. Each subject had peak-to-peak amplitude calculated for the three intensities at 1KHz and the tinnitus frequency. The slope function across the three intensities at each frequency was calculated as the quotient of the Amplitude Dynamic Range (ADR: relative difference of the high intensity and low intensity N1-P2 response amplitude) and Stimulus Dynamic Range (SDR: relative difference of the high and low stimulus intensity).

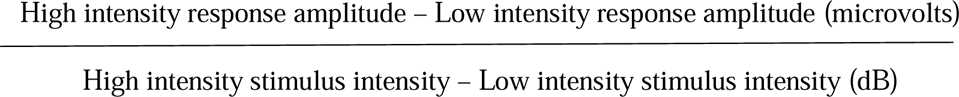

Units of the IDAEP slope are thus μV/dB. Note that the medium intensity stimuli were not used for analysis but were judged important to include in the paradigm for their influence on the overall statistical properties of presented stimuli, and to maintain consistency with other published IDAEP paradigms (Fitzgerald et al., 2009; Carrillo-de-la-Peña et al., 2006).

Participants were rejected from a particular analysis (1 kHz or tinnitus frequency) if they had insufficient overall ERP quality, or less than 5 dB stimulus dynamic range. ERP quality was judged subjectively, and the minimum acceptable SDR was based on inspection of a funnel plot of SDR against IDAEP slope estimate, which showed an inflection point at 5 dB. Such exclusions were performed with the researcher blind to the group to which each participant belonged.

Statistical analysis was performed using the Statistical Package for Social Science (SPSS). Based on the Shapiro-Wilk test of normality, the data were normally distributed across the three groups for 1 kHz and tinnitus frequency for the N1-P2 amplitude, and hence parametric statistics were used. However, the data were normally distributed for 1 kHz and not normally distributed for tinnitus frequency for the slope, and hence a parametric statistic was used for 1 kHz and non-parametric statistic was used for the tinnitus frequency separately.

For the N1-P2 amplitude across groups, a three-way factorial Analysis of Covariance (ANCOVA) was conducted to examine the effects of Group (Acute vs Chronic vs Control), Frequency (1 kHz vs tinnitus frequency), and Intensity (low vs medium vs high) on the N1-P2 amplitude. Stimulus presentation level (which was individualised for each subject) was included as a covariate to control for its potential influence on the dependent measure. Main effects and interactions were assessed, and post hoc comparisons were corrected using Bonferroni test where appropriate. Similarly, a fully repeated measures ANOVA was conducted with Group (Acute vs Post Acute Tinnitus), Frequency (1 kHz and tinnitus frequency), and Intensity (low, medium, high) as within-subject factors. Main effects and interaction effects were tested. Sphericity was assessed using Mauchly’s test, and Greenhouse–Geisser corrections were applied where necessary. Post hoc comparisons were adjusted using the Bonferroni correction.

As the principal analysis of interest, the IDAEP slope values (from N1-P2 response amplitude) for each frequency were compared between the groups. As there was sample size inequality, and a difference in data distribution, between the analyses for each stimulus frequency (Table 2), statistical analysis was conducted independently for each frequency instead of a factorial analysis with group and frequency. Significant differences across the groups for 1 kHz were determined using a one-way ANOVA, and for the tinnitus frequency, the non-parametric Kruskal Wallis test was used. To compare the groups between the Acute phase and the follow-up Post-acute phase for both frequencies, a paired t-test was employed; this constituted the primary dependent variable of interest. We had a very clear prior hypothesis, and therefore used one-tailed statistics to only seek a decrease in IDAEP slope longitudinally over time. For all other analyses, two-tailed statistics were used. Where differences in stimulus presentation intensity or stimulus dynamic range were present, we further carried out an analysis of covariance across groups, treating the stimulus intensities as covariates.

**Table 2:**
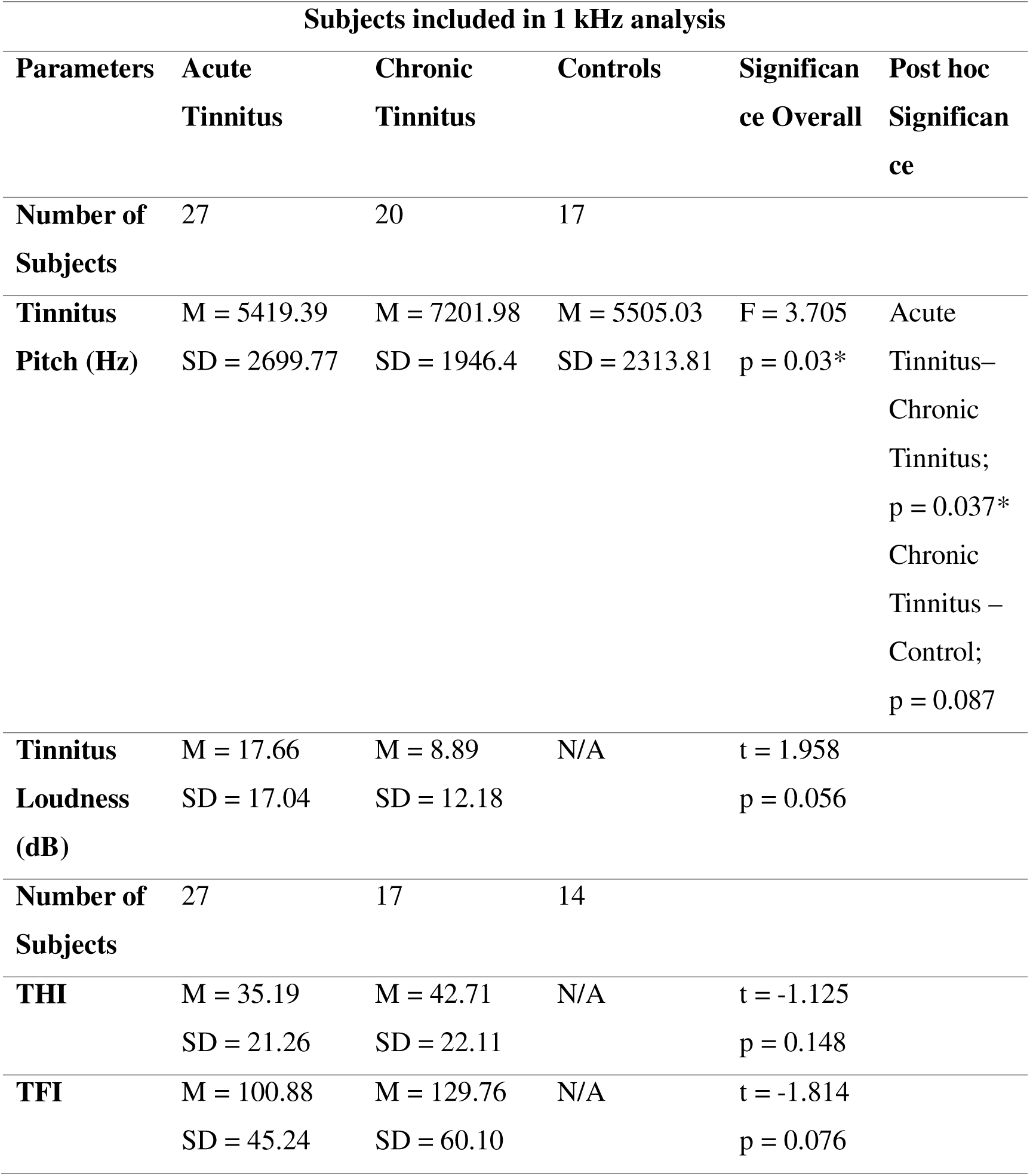
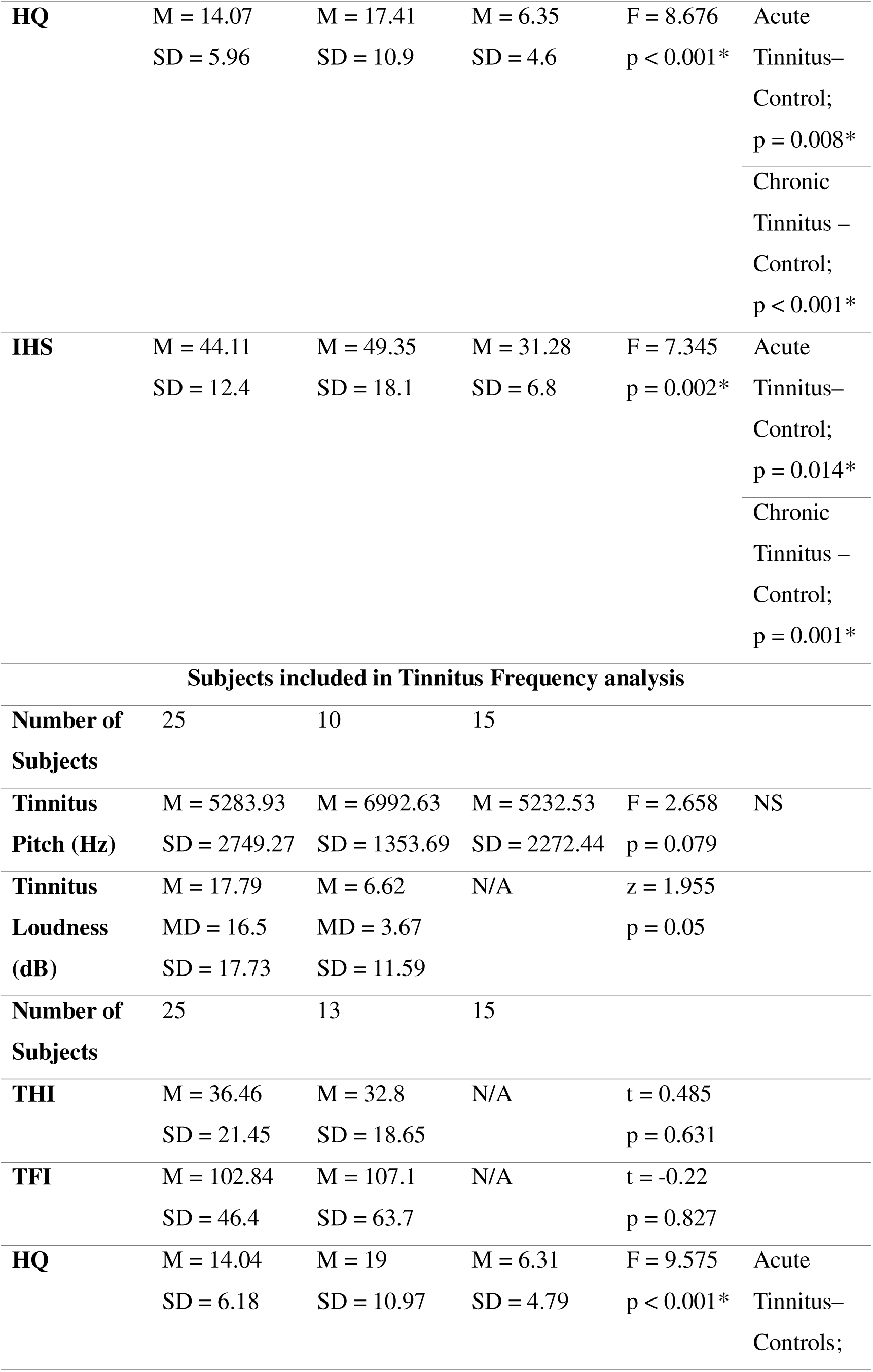
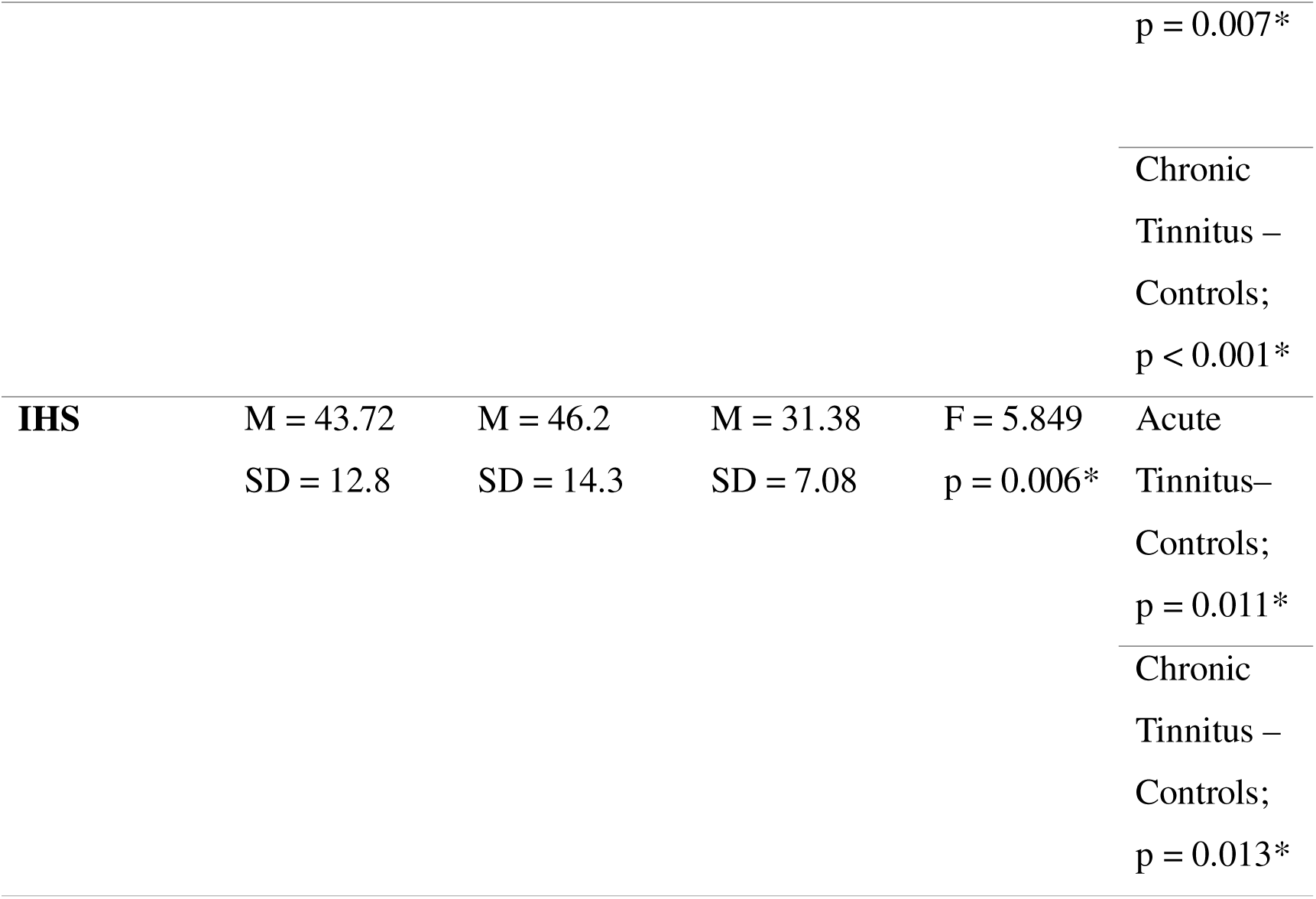
The participants’ tinnitus distress and Tinnitometry related data for each group are shown in Table 2. N denotes number of samples, M denotes mean, SD denotes Standard Deviation, F statistic indicates a One Way ANOVA has been carried out, t statistic indicates an independent t test done, N/A-Not Applicable, NS denotes No Significance. * indicates presence of statistical significance at 95% confidence interval, THI - Tinnitus Handicap Inventory, TFI – Tinnitus Functional Index, HQ – Hyperacusis Questionnaire, IHS – Inventory of Hyperacusis Symptoms. It is important to highlight that there is an asymmetry in the sample sizes between the questionnaires and the original sample, resulting from the exclusion of participant questionnaires with incomplete responses.

As there might be cross-sectional differences and longitudinal changes in tinnitus distress, which might in principle correlate with IDAEP, we also carried out a Pearson correlation between IDAEP slope at 1 kHz and THI score, which we also considered important to exclude any distress-related effects that might otherwise be mistaken for correlates of tinnitus duration. This included one analysis using both Acute and Chronic tinnitus groups combined, and a correlation between the change in IDAEP slope and the change in tinnitus distress from the acute to post-acute phase, reflecting how variations in one over time relate to variations in the other.

Finally, we also conducted an exploratory analysis to investigate the relationship between stimulus dynamic range (SDR) and IDAEP slope, to assess whether differences in SDR might partially explain or mask group differences in IDAEP slope. A further reason for this analysis was to identify a minimum threshold for SDR as an additional quality control measure (see earlier methods relating to setting the minimum acceptable SDR at 5 dB), ensuring that no SDR outliers confounded the results. For each of the three groups, at each stimulus frequency, Pearson’s correlation tests were performed to examine the direction and strength of the association between SDR and slope.

## Results

Final participant groups after data quality and SDR exclusions comprised 27 Acute Tinnitus, 13 Post-acute Tinnitus (Acute Tinnitus individuals followed up six months after tinnitus onset), 20 Chronic Tinnitus, and 17 Controls for the 1 kHz stimulus condition. For the tinnitus frequency condition, the final numbers comprised 25 Acute Tinnitus, 11 Post-acute Tinnitus (Acute Tinnitus individuals followed up six months after tinnitus onset), 13 Chronic Tinnitus, and 15 Controls. The differences in subject group numbers between the two analyses reflects different numbers of instances of individuals’ data meeting exclusion criteria. While estimating the dynamic range for subjects at some of the tinnitus frequencies, there were issues where we could not measure the ULL at high frequencies for the participants due to the limits of sound intensities with our equipment, particularly in participants who had increased degrees of hearing loss at high frequencies. In such instances, ULL was estimated to be maximum stimulation plus 10 dB. Participant group demographics and audiometric characteristics are summarised in Table 1.

### Tinnitometry and symptom questionnaires

With respect to the tinnitometry, there was a non-significant trend between Acute and Chronic Tinnitus for the tinnitus loudness (t(45) = 1.958, p = 0.056) with Acute Tinnitus (M = 17.66, SD = 17.04) having increased tinnitus loudness when compared to Chronic Tinnitus (M = 8.89, SD = 12.18). No significant differences were observed for the tinnitus frequency across groups. With respect to the paired analysis between Acute and Post Acute Tinnitus, there were no significant differences for either tinnitus pitch (t(12) = 0.1, *p* = .888.) or tinnitus loudness (z = −1.165, p = 0.244). Similar results were replicated for the tinnitus frequency with statistical differences in tinnitus loudness and no differences in tinnitus pitch. Refer to Table 1 for further details.

With respect to the symptom questionnaires, for 1 kHz there was a main effect of group for HQ (F(2,56 = 8.676, p < 0.001), with controls (M = 6.35, SD = 4.6) having lower HQ scores than Acute Tinnitus (M = 14.07, SD = 5.96) and Chronic Tinnitus (M = 17.41, SD = 10.9). Comparable results were obtained for IHS (F(2,56 = 7.345, p = 0.002) with controls (M = 31.28, SD = 6.8) having lower IHS scores than Acute Tinnitus (M = 44.11, SD = 12.4) and Chronic Tinnitus (M = 49.35, SD = 18.1). There were no significant differences between Acute and Chronic Tinnitus for THI and TFI. A paired comparison between the Acute and Post-Acute Tinnitus group revealed a significant reduction over time in THI score (t(11) = - 2.549, p = 0.0027), TFI score (t(11) = −4.282, p = 0.013 with Acute Tinnitus having higher THI (M = 24.5, SD = 11.88) and TFI(M = 83.08, SD = 32.62) scores when compared to Post-Acute Tinnitus (THI - M = 17.33, SD = 13.08, TFI – M = 83.08, SD = 30.63. For hyperacusis questionnaires, no change over time was observed for scores on either HQ (t(12) = 0.04, p = 0.966) and IHS (t(12) = −0.326, p = 0.75).

Similar results were replicated at tinnitus frequency with an effect for HQ and IHS and no differences between Acute and Chronic Tinnitus for both the tinnitus distress questionnaires (THI and TFI). The results of the symptom questionnaires and tinnitometry for each of the three groups are shown in Table 2.

### N1-P2 amplitude

For cross sectional comparisons between Acute, Chronic, and Controls, after adjusting for stimulus presentation level, the repeated measures ANCOVA revealed significant main effects of Frequency (F(1,332) = 105.24, p < 0.001) with 1 kHz (M = 10.74, SE = 0.264) having higher amplitude than tinnitus frequency (M = 6.59, SE = 0.295) and Intensity (F(2,332) = 4.08, p = 0.018), with high intensity (M = 9.33, SE = 0.36) having higher amplitude than low intensity (M = 7.83, SE = 0.36), but no main effect of group (F(2,332) = 1.929, p = 0.147.) There was no three-way interaction between Group, Frequency, and Intensity (F(2,332) = 0.071, p = 0.991).

With respect to the longitudinal comparisons between Acute and Post Acute Tinnitus, repeated measures ANOVA was carried out and the sphericity was not violated as Mauchly’s Test of Sphericity yielded no significant difference for both simple main effect and interaction effects. Results revealed significant main effects of Frequency (F(1,10) = 19.676, p = 0.001), with 1 kHz (M = 11.75, SE = 1.024) having higher amplitude than tinnitus frequency (M = 5.88, SE = 0.52) and Intensity (F(2,9) = 11.99 p = 0.003) with high intensity (M = 9.62, SE = 0.55) having higher amplitude than low intensity (M = 7.96, SE = 0.46). A significant interaction was observed for frequency and intensity (F(2,9) = 4.607 p = 0.042) with both frequencies significantly differing under each intensity. The three-way Frequency × Intensity × Group interaction (F(2,9) = 1.169, p = 0.354) yielded no significant effect. Figure 2 illustrates the N1-P2 waveforms across intensities and frequencies, and Figure 3 depicts the N1-P2 amplitude across intensities and frequencies.

**Figure 1:**
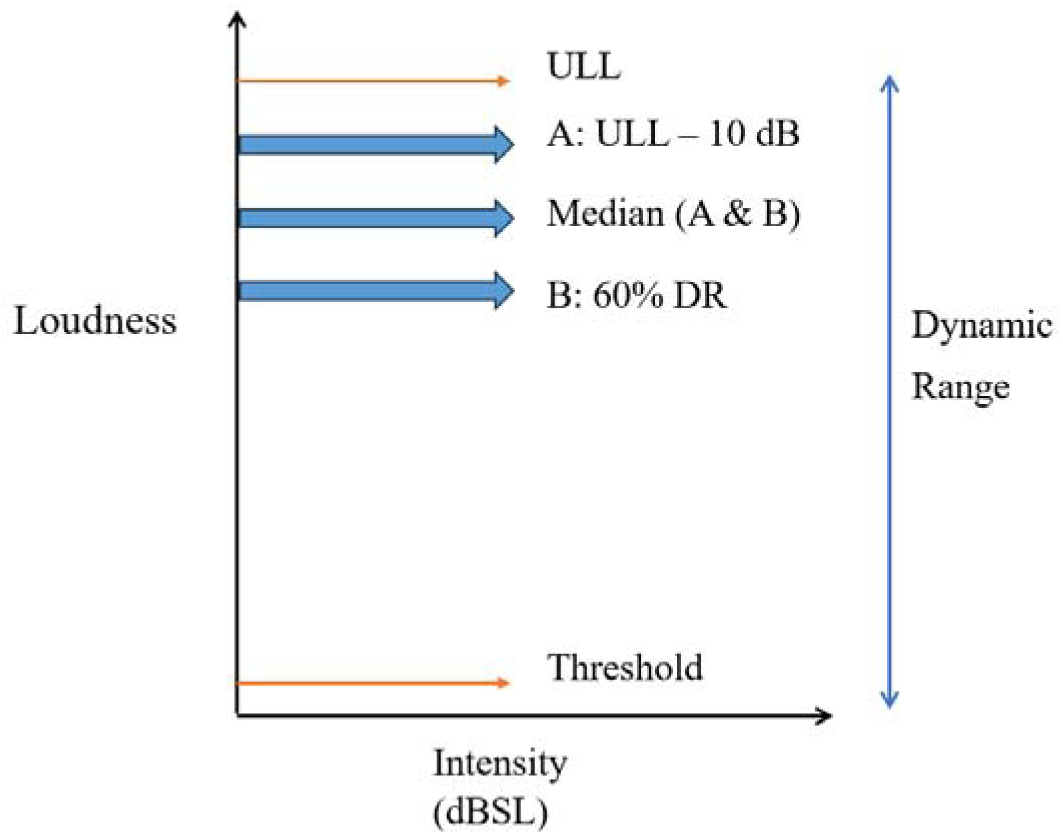
Stimulus presentation levels across 3 intensities based on tailored dynamic range for each subject.

**Figure 2:**
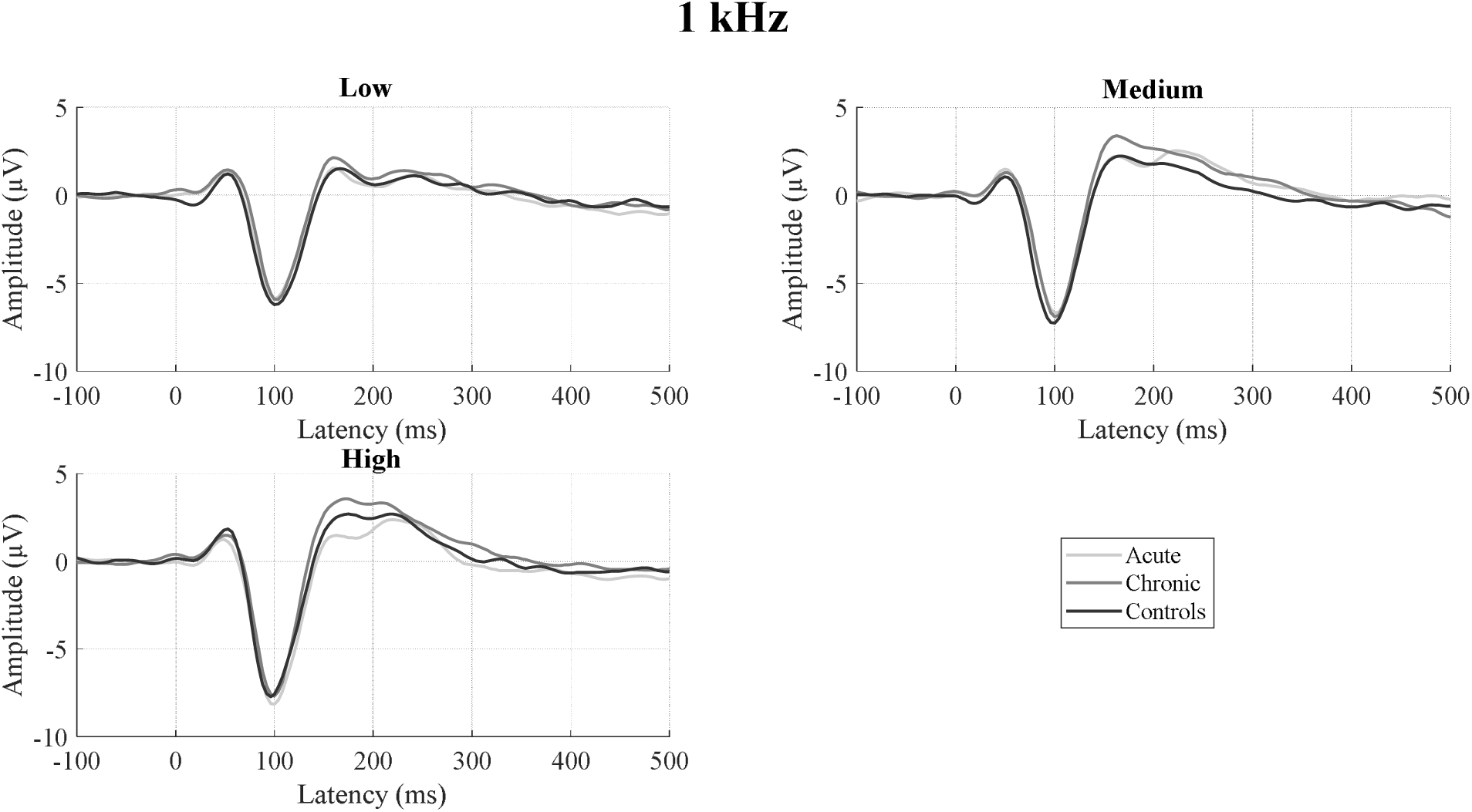
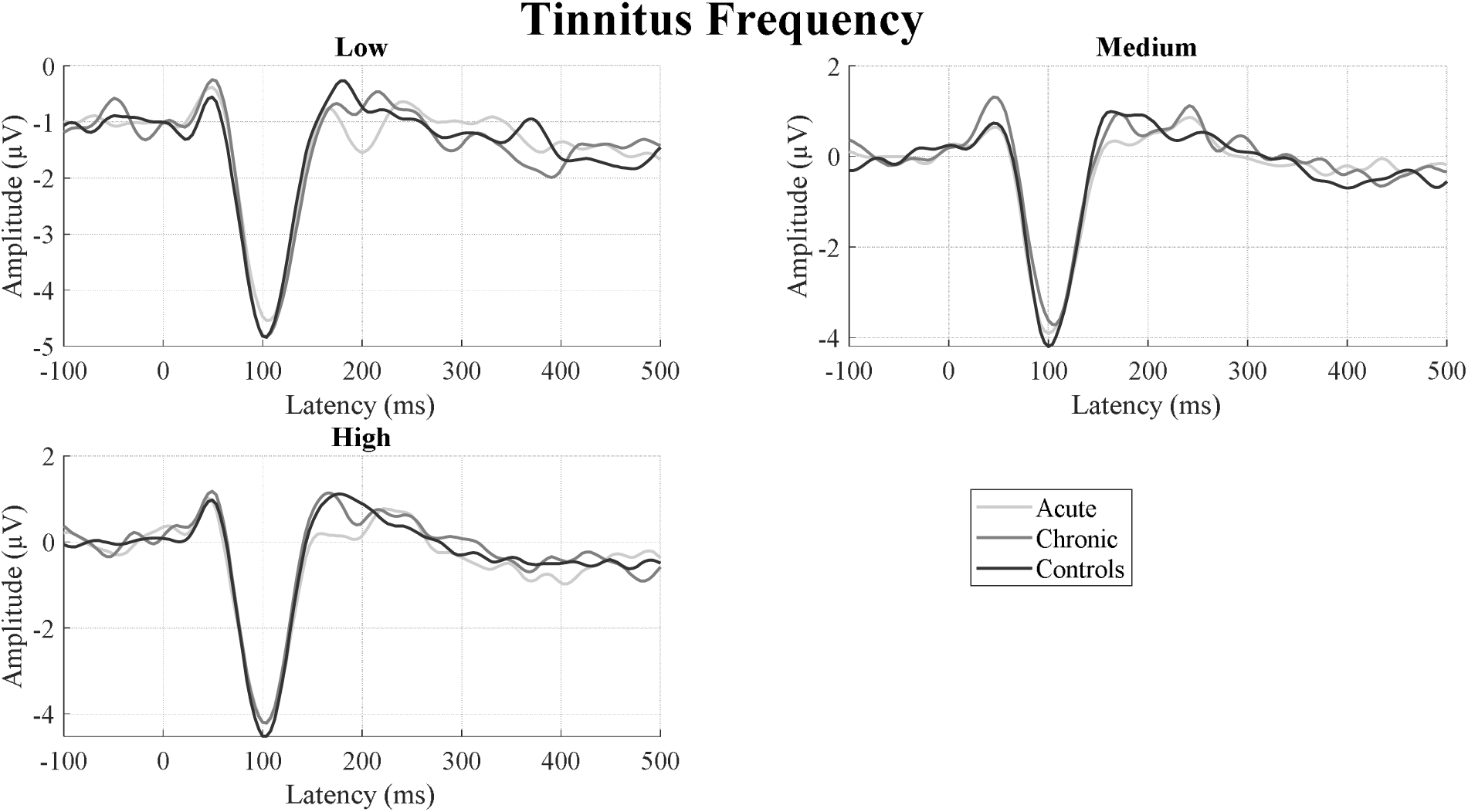
IDAEP waveforms across three intensity levels and two frequencies.

**Figure 3:**
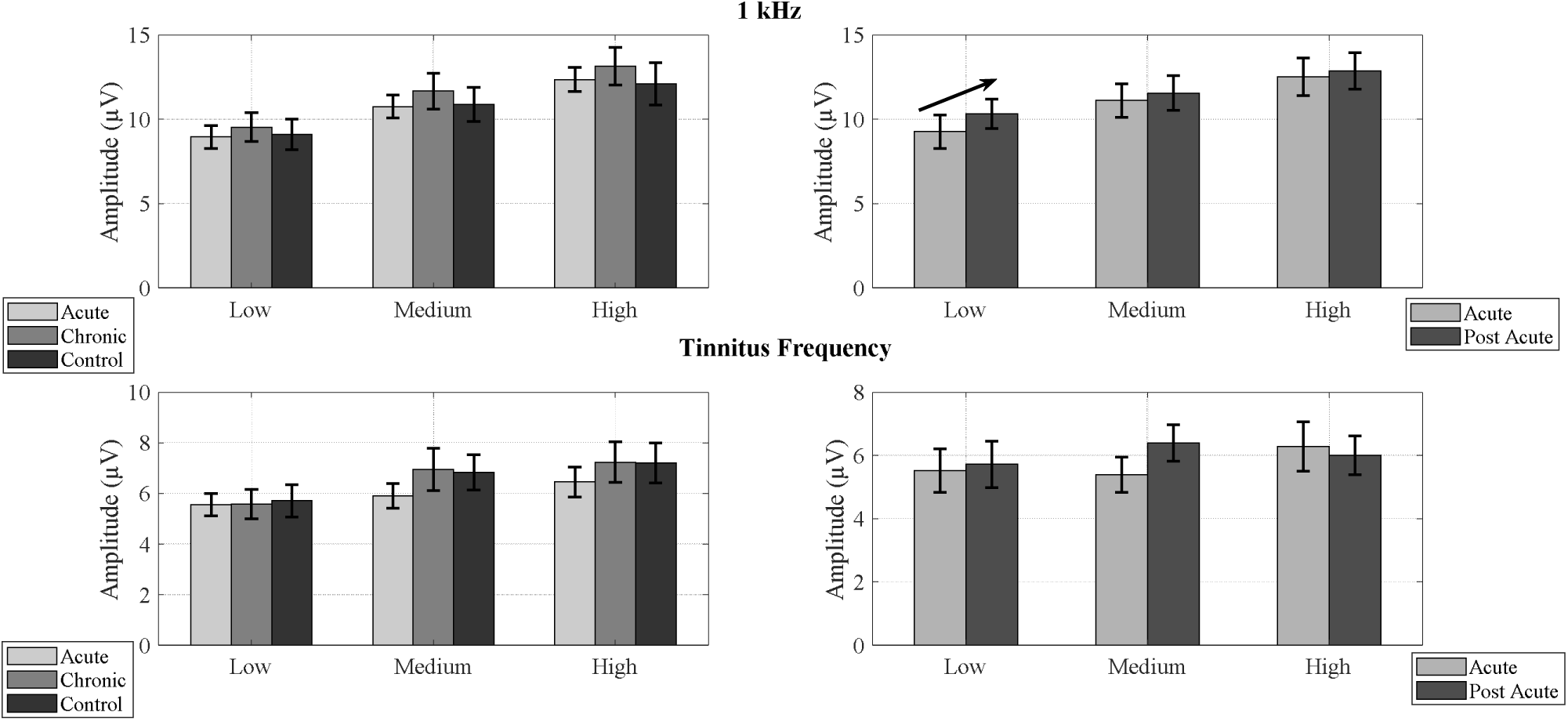
Variations in N1-P2 amplitude between groups and across different stimulus presentation and frequencies. Error bars indicate standard error for mean. Only a non-significant trend was observed between Acute and Post Acute Tinnitus for the low intensity presentation at 1 kHz which is represented by an increasing arrow between the groups. Remaining comparisons yielded no significant differences.

### IDAEP slope

There was no significant effect of group on IDAEP slope comparing the Acute, Chronic, and Control groups cross-sectionally for 1 kHz (F (2,61) = 2.194, p = 0.12) upon one way ANOVA, or for tinnitus frequency (*X^2^*(2) = 2.43, p = 0.297) upon a Kruskal Wallis test. For longitudinal comparisons, a one-tailed t-test comparing changes between Acute and Post-acute stages (only assessing decreases over time) revealed a significant longitudinal decrease in IDAEP slope at 1 kHz over time (t (12) = −1.919, p = 0.04) with the Acute Tinnitus group (M = 0.186, SD = 0.114) having larger slopes than the Post-acute group (M = 0.133 SD = 0.142). Both Acute and Chronic groups had larger IDAEP slopes than Controls, which we mention only to contextualise the longitudinal changes described below, as these cross-sectional differences were not statistically significant. No statistically significant differences were obtained longitudinally between Acute and Post Acute Tinnitus at the tinnitus frequency, though the non-significant changes were in the same direction as for 1 kHz. Figure 4 illustrates the IDAEP slope across groups and frequencies.

**Figure 4:**
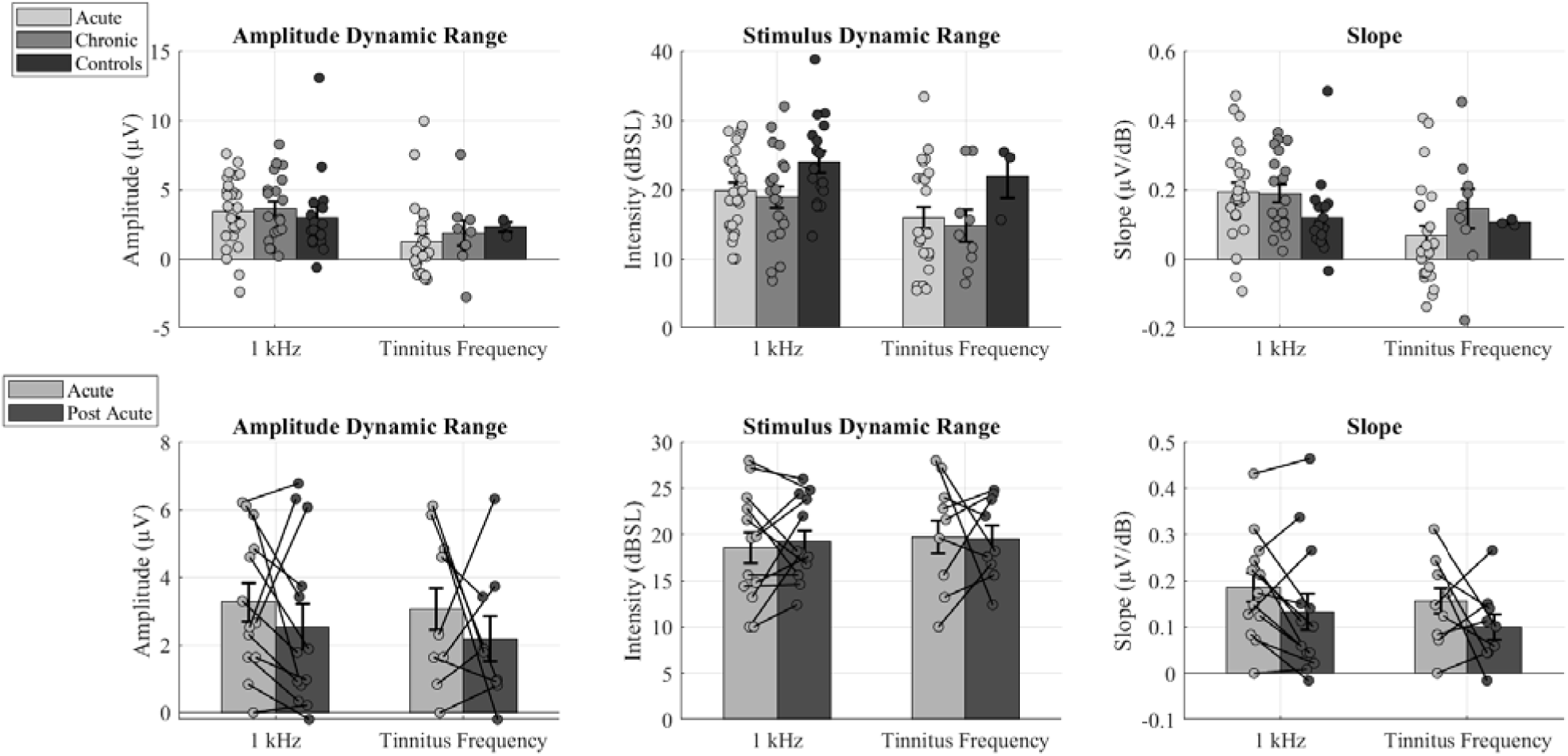
Variations in Amplitude Dynamic Range, Stimulus Dynamic Range, and IDAEP Slope (Amplitude Dynamic Range/Stimulus Dynamic Range) between groups across frequencies. Error bars indicate standard error for mean and TF indicates Tinnitus Frequency. A statistically significant difference was found between Controls and Acute Tinnitus, Controls and Chronic Tinnitus for Stimulus Dynamic Range at 1 kHz. A statistically significant difference was noted between Acute and Post Acute Tinnitus for Slope.

### Stimulus and amplitude dynamic ranges individually

Whilst our primary outcome measure was IDAEP slope (i.e. amplitude dynamic range divided by stimulus dynamic range), we analysed these dynamic range variables separately also.

There were however statistically significant effects of group on the stimulus dynamic range for 1 kHz (F(2,61) = 3.346, p = 0.042). Due to differences in stimulus dynamic range, we carried out an analysis of covariance for IDAEP slope with stimulus dynamic range as a covariate and found no significant effect when comparing the Acute, Chronic, and Control groups cross-sectionally (F (2,60) = 1.608, p = 0.209).

Longitudinally, the group mean stimulus dynamic range did not change between Acute and Post-acute stages (t(12) = 0.482, p = 0.639; 1 kHz, t(10) = 0.036, p = 0.972; tinnitus frequency), and there was a non-significant trend in the amplitude dynamic range (t(12) = - 1.099, p = 0.293; 1 kHz, t(10) = −1.873, p = 0.09; tinnitus frequency)

### Relationship between Tinnitus Distress and IDAEP slope

Considering the presence of statistically significant differences between Acute and Post-acute Tinnitus longitudinally for both tinnitus distress (THI and TFI) and IDAEP slope at 1 kHz, we explored the relationship between tinnitus distress (THI) and IDAEP slope. A Pearson correlation analysis cross-sectionally between tinnitus distress and IDAEP slope including both Acute and Chronic Tinnitus participants revealed a statistically significant negative correlation (*r*(41) = −.356, *p* = .019). However, a similar correlation between the Acute-to-Post-acute change in IDAEP slope and the Acute-to-Post-acute change in tinnitus distress was not observed (*r*(9) = .185, *p* = .586). Figure 5 illustrates the changes between tinnitus distress and IDAEP slope.

**Figure 5:**
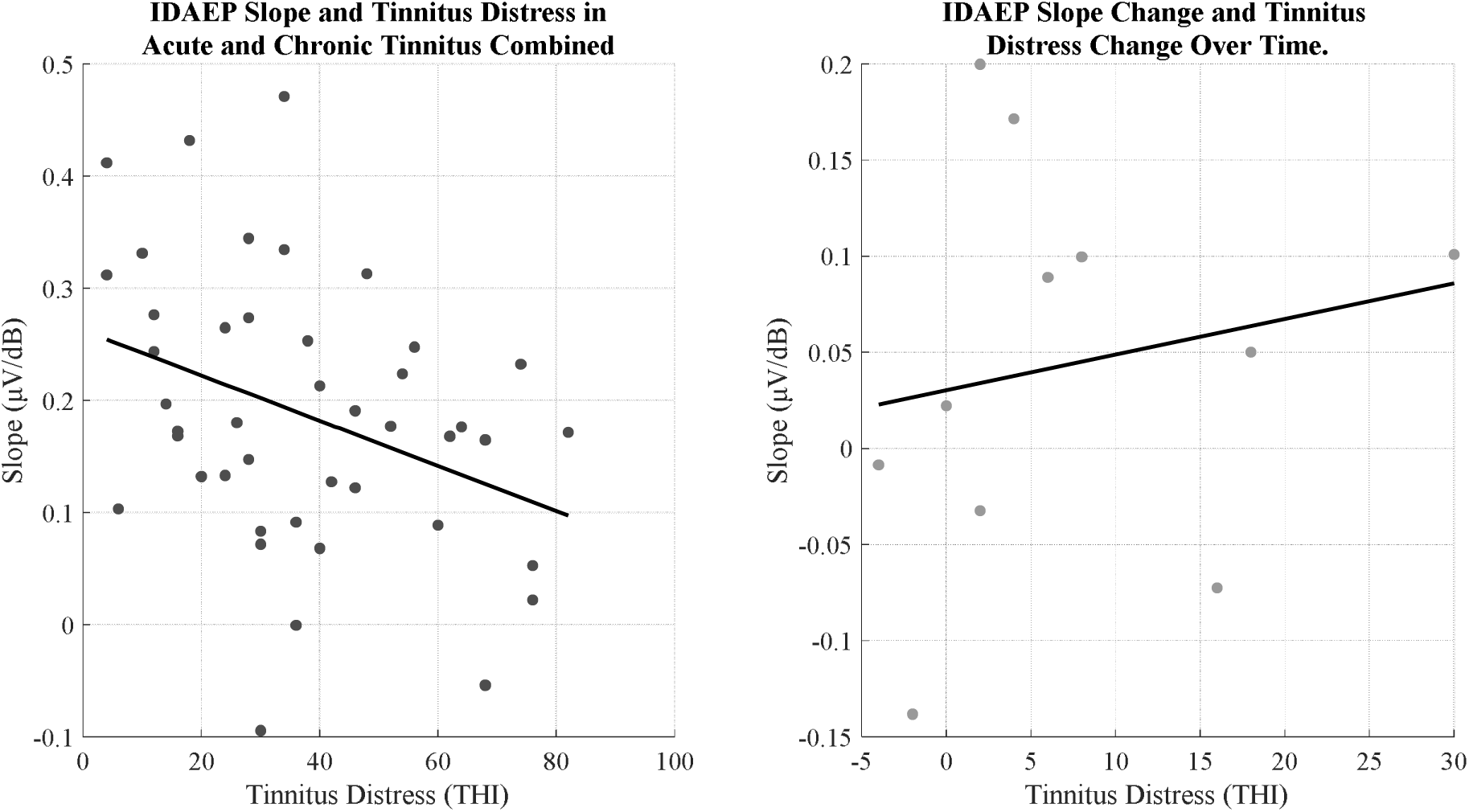
Relationship between Tinnitus Distress and IDAEP slope across both Acute-Chronic and Acute-Post Acute groups. THI indicates Tinnitus Handicap Inventory and there is a presence of statistically significant negative correlation between tinnitus distress and IDAEP slope across Acute-Chronic Tinnitus groups.

### Relationship between IDAEP slope and stimulus dynamic range

An additional exploratory analysis was conducted for the 1 kHz and tinnitus stimulus frequencies to determine the association between IDAEP slope and stimulus dynamic range (SDR). This was partly as a control analysis, to ensure that any differences in IDAEP slope between groups were not due to differences in stimulus dynamic range between groups, on account of dynamic range adaptation. Furthermore, we were interested to determine whether different stages of tinnitus were associated with differences in dynamic range adaptation for IDAEP, which might manifest as different relationships between dynamic range and IDAEP slope.

A non-significant trend of a negative correlation was found for the Acute tinnitus group (Correlation: r(25) = −0.356, β = −.0081, *p* = .068), and no correlation was observed for the chronic tinnitus group (Correlation: r(18) = 0.086, regression: β = .0014, *p* = .717), or for the non-tinnitus controls (Correlation: r(15) = 0.156, β = .0032, *p* = .485). There was no significant correlation for all groups across tinnitus frequency and no significant correlations in both Acute and Post Acute Tinnitus for 1 kHz and tinnitus frequency (all p-values > 0.05). For further details, please refer to Figure 6.

**Figure 6:**
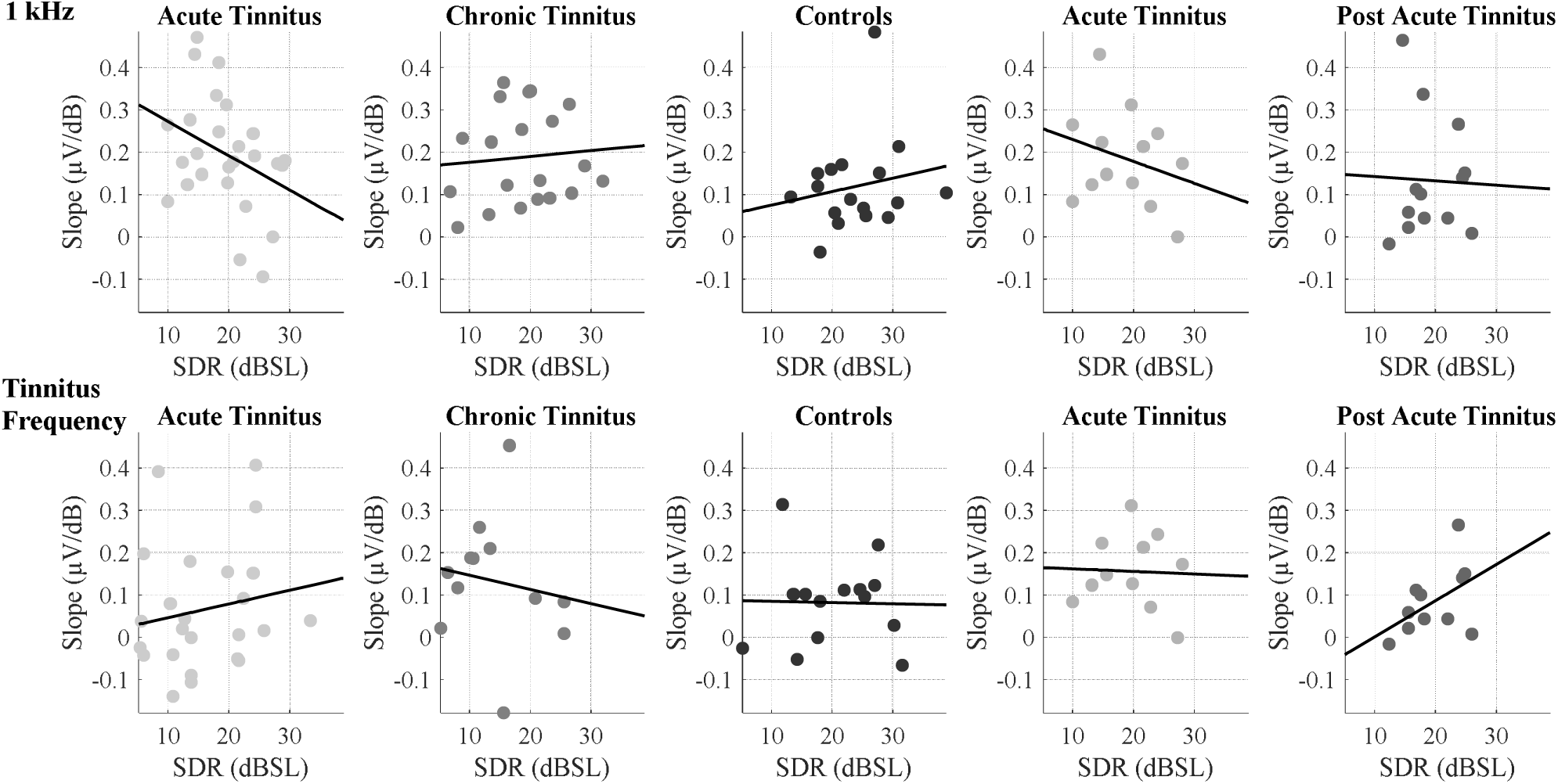
Represents the relationship between slope and stimulus dynamic range across groups and frequencies. SDR indicates Stimulus Dynamic Range. A non-significant negative trend was noted for Acute Tinnitus at 1 kHz.

## Discussion

### IDAEP is increased in the acute stages of tinnitus

Our hypothesis postulated an increase in auditory hyperexcitability and/or hypervigilance associated with the onset of tinnitus, acting in combination with mechanisms compensating for reduced auditory input, effectively increasing the precision of auditory inputs. Together, we hypothesised that these processes lead to perception of spontaneous auditory pathway activity as a sound source. As central plastic processes governing memory and predictions relating to the tinnitus percept become engaged, we also hypothesised that the hyperexcitability/hypervigilance state surrounding tinnitus onset need only be transient for chronic tinnitus to occur, and as such that we would see initial elevation of these markers around tinnitus onset, and subsequent reduction for the majority of the group reflecting regression to the mean (Sedley et al., 2016). Our results of increased IDAEP slope at 1 kHz in Acute Tinnitus individuals that further reduces over time during the Post-acute stages of tinnitus aligns with this hypothesis. With respect to the sensory precision model, heightened IDAEP slope may represent a specific type of auditory hypervigilance (or precision control mechanism) relevant to the tinnitus signal.

Although there is no prior research into acute tinnitus and event related potentials, our recent research on subjective sensitivity changes between acute and post-acute tinnitus (including some of the same subjects featured in the present study) (Umashankar et al., 2025) has found that another potential marker of central gain (subjective auditory sensitivity, in the form of loudness growth slope and hyperacusis symptom questionnaires) does not vary between acute and post-acute tinnitus. This distinction highlights the importance of distinguishing specific aspects and indicators of gain, and their individual relationships to tinnitus.

### IDAEP and (tinnitus) distress

As an alternative or additional possibility, we considered whether IDAEP reflects the degree of distress associated with tinnitus, because both tinnitus distress (THI and TFI) and IDAEP slope have now been found to decrease over time between in Post Acute Tinnitus. IDAEP changes might have simply been explicable as correlates of (changing) tinnitus distress. However, our findings of no correlation between tinnitus distress and IDAEP slope across the longitudinal data of Acute and Post Acute Tinnitus, makes this unlikely to be an explanation for the longitudinal changes in IDAEP at a group level. To the contrary, our findings, within the combined group of Acute and Chronic Tinnitus subjects, of a significant *negative* correlation between IDAEP slope and tinnitus-related distress is in the opposite direction, and somewhat unexpected. We are cautious to avoid overinterpretation of this correlation result in isolation, especially as it was an exploratory analysis. Although literature on tinnitus and IDAEP is limited, prior studies in depressive disorders also report *increased* IDAEP in affected individuals (Lee et al., 2014; Kangas et al., 2024). This divergence highlights the need for further investigation, particularly into the directionality of IDAEP changes in relation to tinnitus distress. Longitudinal studies, especially in individuals with acute tinnitus and escalating distress, might help to elucidate these dynamics over time (beyond 6 months), and would complement this study, in which the group showed a reduction in distress over time.

### Relative lack of IDAEP differences at high frequencies

The tinnitus frequency did not exhibit a strong mean IDAEP slope, and in some subjects, it was zero or negative. This could in part be because of the individuals’ lower stimulus dynamic ranges at the tinnitus frequency, due to the presence of hearing loss at high frequencies in all our groups. Due to the limits of sound intensities possible with our equipment at high frequencies affected by hearing loss, there were some subjects in whom we could not determine their ULL at the tinnitus frequency; stimulus frequency in these subjects was thus limited by technical possibility rather than ULL. Finally, as IDAEP to high frequency stimuli is not well-studied in the literature, it may simply be the case that the N1-P2 complex at high frequencies is less intensity-dependent. However, it is notable that the non-significant trends observed, cross-sectionally and longitudinally, were in keeping with results observed for 1 kHz stimuli.

### Need to explore Serotonergic variations in tinnitus

The IDAEP has also been validated as a reliable tool for assessing serotonergic function, showing a positive correlation with serotonin levels. A steep IDAEP slope suggests reduced serotonin levels or action, and increased hypervigilance or hyperreactivity to sensory stimuli, whereas a shallow slope indicates the opposite (Carrillo-de-la-Peña et al., 2006). Recent PET studies further support a link between the IDAEP slope and central serotonergic activity, particularly in the temporal cortex (Pillai et al., 2020), reinforcing its potential as a marker for serotonin variations. There has been literature reporting the influence of serotonergic variations in tinnitus (Sun et al., 2007; Simpson & Davies, 2000). Tang & Trussell (2015) suggested that serotonin significantly alters the intrinsic properties of fusiform cells, potentially increasing the sensitivity of the dorsal cochlear nucleus to sensory input and contributing to the generation of tinnitus. A post-mortem study revealed a significant reduction in serotonergic neurons within the dorsal and obscurus raphe nuclei of tinnitus patients compared to non-tinnitus patients, highlighting the potential involvement of the brainstem serotonergic system in tinnitus pathology (Almasabi et al., 2022). There is, to our knowledge, no prior published work relating serotoninergic function to acute tinnitus. The wider literature on serotonin and tinnitus focuses on chronic tinnitus and often on measuring serotonin levels in whole blood rather than directly within the central nervous system (Sachanska, 1999). The findings from these studies are inconsistent, while some report elevated serotonin levels in tinnitus patients (Liu et al., 2003; Clewes, 2012) other studies suggest a serotonergic deficiency in tinnitus (Almasabi et al., 2022). Our IDAEP findings of a decrease in slope from the Acute to the Post-acute stage of tinnitus may indicate a hyposerotonergic state during the Acute phase of tinnitus. We propose this as a hypothesis for future research, which might have substantial implications for mechanisms of tinnitus onset and/or early specific treatment opportunities.

### Dynamic range adaptation in acute tinnitus warrants future study

It has been argued that tinnitus may be at least partly independent and distinct from central gain, a commonly proposed mechanism for tinnitus and hyperacusis pathophysiology (Sedley et al., 2016; Knipper et al., 2013; Zeng, 2013). Central gain is essentially a vertical scaling of a neuron’s input output function and may be argued to relate more to hyperacusis than tinnitus (Zeng, 2013). Our exploratory investigation of the relationship between IDAEP slope and stimulus dynamic range suggests there might be a more nuanced relationship to tinnitus; the onset period of tinnitus may be associated with increased dynamic range adaptation, which can be considered a horizontal scaling or shift of a neuronal system’s input-output function to match the distribution of input stimulus intensities. We observed a non-significant trend of negative correlation between IDAEP slope and stimulus dynamic range, which was specific to the Acute tinnitus group. This pattern is consistent with dynamic range adaptation, whereby neurons respond more sensitively (i.e. show greater IDAEP) to smaller dynamic, and less sensitively (i.e. lesser IDAEP) to larger stimulus dynamic ranges. This preliminary finding raises the possibility of an increase in dynamic range adaptation in acute tinnitus, which, if confirmed in future studies, could indicate enhanced adaptive mechanisms associated with the onset of this condition. Consistent with this possibility, Diesch & Hassel-Adwan (2017) explored dynamic range adaptation in their study, where they used a varied amplitude modulation tones in both ascending and descending run to compare tinnitus with controls. They found that tinnitus individuals had stronger undershoot (strong decrease in amplitude with decrease in stimulus) and weaker overshoot (small increase in amplitude with increase in stimulus). In keeping with this, Yukhnovich et al. (2024) also observed that, once hyperacusis had been excluded, tinnitus was associated with larger mismatch negativity (MMN) responses to downward intensity deviants, which was best explained by an altered state of adaptation. Very limited literature has directly searched for the presence of dynamic range adaptation in tinnitus, and our tentative results indicate that this area should further be explored. However, as the finding was non-significant, we present this as simply a hypothesis for future study rather than a conclusive finding. An additional limitation of our findings is that the multiple dynamic ranges were not contrasted within individuals with each individual needed to be presented with stimulus sets from multiple dynamic ranges.

### Conclusion

We report the novel finding that during acute tinnitus, IDAEP, often regarded as a correlate of hypervigilance and central gain, and with links to central serotonergic function, tends to increase around the acute stage, and then return toward normal during the chronic stages, of tinnitus. Future research should consider the onset conditions of tinnitus, as well as its enduring long-term correlates, which we have demonstrated here can look very different. Future studies should also specifically assess central serotonergic function and dynamic range adaptation in neuroimaging and IDAEP paradigms respectively, to further elucidate the mechanism of tinnitus generation and persistence.

